# TRPV2 activation reorganizes actin cytoskeleton, induces neurite initiation and branching by altering cAMP levels

**DOI:** 10.1101/817684

**Authors:** Manoj Yadav, Chandan Goswami

## Abstract

Understanding of molecules and their role in neurite initiation and/or extension is not only helpful to prevent different neurodegenerative diseases but also can be important by which neuronal damages can be repaired. In this work we explored the role of TRPV2, a non-selective cation channel in the context of neurite functions. Using western blot, immunofluorescence, and live cell Ca^2+^-imaging; we confirm that functional TRPV2 is endogenously present in the F11 cell, a model system mimicking peripheral neuron. In F11 cells TRPV2 localizes in specific sub-cellular regions enriched with filamentous actin, such as in growth cone, filopodia, lamellipodia and in neurites. TRPV2 regulates actin cytoskeleton and also interacts with soluble actin. Ectopic expression of TRPV2-GFP but not GFP-only in F11 cell induces more primary and secondary neurites, confirming its role in neurite initiation, extension and branching events. TRPV2-mediated neuritogenesis is dependent on wild-type TRPV2 as cells expressing TRPV2 mutants reveal no neuritogenesis. However, TRPV2-mediated neuritogenesis is unperturbed by the chelation of intracellular Ca^2+^ by BAPTA-AM, and thus involves Ca^2+^-independent signaling events also. We demonstrate that pharmacological modulation of TRPV2 alters cellular cAMP levels. These findings are relevant to understand the sprouting of new neurites, neuroregeneration and neuronal plasticity at the cellular, subcellular and molecular level. Such understanding may have border implication in neurodegeneration and peripheral neuropathy.

## Introduction

Regeneration of neurites, especially in the peripheral tissue has immense importance in the context of different neuronal disorders. Importance of Ca^2+^-signaling and other signaling pathways in the context of neuritogenesis is well established, yet the molecular mechanisms and the molecular candidates involved in these processes are poorly understood^1–7^. Neuritogenesis is a complex process by which new neurites are originated from cell body followed by series of stochastic events such as neurite extension and/or retraction, bending and often branching at certain points^2, 7–8^. Regulation of membrane proteins and membrane dynamics, vesicular recycling, sub-membranous cytoskeleton and subsequently gross cytoskeletal reorganization are major events involved in these processes. All these events are primarily regulated by an array of regulatory proteins that are present on the cell surface, such as ion channels, receptors and adhesion molecules which sense the different chemical signaling cues and allow the neurons to respond accordingly^8–10^. It is well established that inhomogeneous distribution of Ca^2+^-levels is a crucial parameter for most of these processes^2^. Indeed, changes in the spatiotemporal Ca^2+^-levels and Ca^2+^-oscillation patterns have been correlated with most of these functions^2, 8–10^. Such aspects strongly suggest the importance of different Ca^2+^ channels in the regulation of different aspects of neuritogenesis. Neuritogenesis process is critical for proper contact formation, synapse formation and thus for proper neuronal functions. Therefore, understanding the neuritogenesis process at cellular and molecular level is relevant to several neurological disorders including several forms of peripheral neuropathy and neurodegeneration.

In this context previous studies have shown the involvement of TRPV channels, Ca^2+^ and other signaling molecules as an intermediate to regulate this complex process. Such as rapid dynamics of neurite extension or retraction in response to chemical cues is well established^11^. Ca^2+^-dynamics and thus different Ca^2+^ channels are known to regulate the neuronal growth cone dynamics^12^. Rapid neurite outgrowth can also be triggered by laser-based micro-heating, suggesting that neurites have unique ability to sense minor temperature differences also^13^. Similarly, mechanosensitive Ca^2+^-channels have also been implicated in the initiation of neuritogenesis^14^. Transient Receptor Potential Vanilloid channels (TRPVs) are members of non-selective cation channels, most of these channels have precise thermosensitive and mechanosensitive behavior, are known to be expressed in peripheral neurons and are involved in several neuronal functions^15–19^. Endogenous expression of different TRPVs in specific neurons and in a specific region of neuronal tissues are highly suggestive of their precise role in neuronal contact formation. Indeed, the importance of TRPV members, particularly for the TRPV1 and TRPV4 in the context of neurite and synaptic functions have been reported^8–9, 20–24^. Other than TRPVs, few other TRP channels and Ca^2+^ channels have also been implicated in such functions. For examples, TRPC5 has been implicated in the regulation of neurite movement and growth cone morphology^25^. TRPC channels has also been implicated in the netrin-1-induced chemotropic turning of nerve growth cones^26^. In spite of all these reports, full array of Ca^2+^-channels involved in these process have not been identified and the exact molecular and cellular events involved in such events are not common and not well-understood.

In this regard we seek to understand the mechanism of neuritogenesis process and its key signaling partners. Based on the existing studies and expression profile we hypothesize that TRPV2 plays important role in the neuritogenesis process and it is regulated by the cAMP level in the growing neurites. It also plays an important role in different sensory functions. Recently involvement of TRPV2 in the regulation of neurite has been demonstrated^10, 27^. TRPV2 knockout animals show several abnormalities like cardiac dysfunction and altered immune response^28–29^. Such altered-functions can be attributed to the altered neuronal circuit formation and neuro-immune interaction^30–32^. TRPV2 plays an important role during neuronal contact formation and also in the process of synaptic plasticity, though the molecular mechanisms involved in such process have not been investigated. Therefore, in this work, we have explored the importance of TRPV2 in specific neuronal functions such as neurite initiation, neurite extension and neurite branching by using confocal imaging and biochemical tools.

## Materials and methods

### Reagents and antibodies

Probenecid (P8761), Tranilast (T0318) were purchased from (Sigma-Aldrich, St Louis, MO, USA). All secondary IgG antibodies (alexa-488-labelled anti-mouse (A-21200), alexa-488-labelled anti-rabbit (A-11070), alexa-594-labelled anti-rabbit (A-21442), alexa-594-labelled anti-rat (A-21472), DAPI (D1306), Alexa-488-labelled Phalloidin (A-12379) and Ca^2+^-sensor dye Fluo4-AM (F14201) were purchased from Invitrogen (Carlsbad, California, USA). Fluoromount-G (0100-01) was purchased from Southern Biotech (Birmingham, US). Anti TRPV2 antibody (Rabbit ACC-039) for an extracellular loop was purchased from Alomone lab (Jerusalem, Israel). Detailed list of all the reagents used in the study have been provided in supplementary file.

### Cell culture, transfection and DNA constructs

F11 and CHO K1 cells were cultured in F12 Hams medium (Himedia AL025) supplemented with 10% FBS (Himedia RM9970), 2mM L-glutamine (Himedia TCL012), penicillin-streptomycin (Himedia A018) (100 units/ml), amphotericin-B (sigma A2942) 2.5ng/μl. HaCaT, HEK 293, SaOS and Neuro2a cells were cultured in DMEM medium (Himedia) supplemented with 10% FBS (Himedia, RM9970), 2mM L-glutamine (Himedia), penicillin-streptomycin (Himedia), (100 units/ml), amphotericin-B (sigma) 2.5ng/μl. Cells were maintained in a humidified atmosphere at 5% CO_2_ and 37°C. All cells were maintained in a humidified atmosphere at 5% CO_2_ and 37°C. For transient transfection, Lipofectamine 2000 (11668027) and Lipofectamine 3000 (L3000015) (Invitrogen) was used according to the manufacturer’s instructions. For transient expression of TRPV2, fluorescent protein-tagged, plasmid encoding TRPV2-Wt-GFP, TRPV2-N571T, TRPV2-N572T, TRPV2-NN571-572TT, and actin-RFP were used^22, 33^. Approximately 30 hours after transfection the cells were fixed by PFA (Sigma). GFP-only was expressed by using pEGFPN3 vector. cAMP sensor Epac-S^H188^ (mTurq2Δ_Epac (CD, ΔDEP) td ^cp173^Ven) was a kind gift from Prof. Kees Jalink^34^.

### Immunofluorescence analysis

For immunocytochemical analysis cells were fixed with 4% PFA for 5 min. Fixed cells were permeabilized with 0.1% Triton X-100 (T8787) in PBS (5 min) and subsequently blocked with 5% BSA (BSASG100). Primary antibodies raised against TRPV2 was used in 1:500 dilution. Alexa 488-conjugated secondary antibody was used at 1:1000 dilution.

### BAPTA-AM treatment

To determine the effect of BAPTA-AM on F11 cells. F11 cells were treated with 10 μM BAPTA-AM and were either treated with TRPV2 activator (Probenecid) or TRPV2 inhibitor (Tranilast). To see the cellular processes and morphology cells were stained with Phalloidin, Microtubules and DAPI.

### Flow cytometry

To check the level of tyrosine kinase, CREB and P-CREB after long term activation cells were stained with specific antibody mentioned above. Followed by flow cytometry analysis. Stained cells were washed three time with the FACS buffer prior to acquisition. Acquisition was performed with FACS Calibur (BD Biosciences). Analysis of data was performed by using CELL QUEST PRO software (BD Biosciences).

### Ca^2+^-imaging

F11 cell were cultured on a 25mm glass coverslip. Around 24 hours after seeding, cells were loaded with Ca^2+^-sensitive dye (Fluo-4 AM, 2 μM for 30 min). Subsequently, cells were placed in the live cell chamber and images were acquired. Cells were stimulated with specific agonists or antagonists at the specific frame as described. Fluo-4 AM signal was acquired using NIKON confocal microscope. The images were analyzed using Fiji software and intensities specific for Ca^2+^-loaded Fluo-4 are represented in artificial rainbow color with a pseudo scale (red and blue indicating the highest and lowest levels of Ca^2+^ respectively). For experiments aiming at chelation of intracellular Ca^2+^, BAPTA-AM (Sigma A1076) was used at 10microM. In such experiments with minor modification of the protocol described before^8^. Cells were seeded in the optimum density (60%) on glass cover slip. Approximately 12 hours after seeding BAPTA-AM was added on the cells and 3hrs after BAPTA-AM treatment, TRPV2 specific agonist and antagonist were added on the cells. After 16 hours of drug treatment, the cells were fixed with 4% PFA.

### Live cell imaging

F11 cell were cultured on 25mm glass cover slip in complete media. Cells were imaged 36 hours after transfection with TRPV2-GFP and/or Actin-RFP. Cells were imaged with ZEISS LSM-780. In some experiments aimed to evaluate the effect of TRPV2 modulation, certain pharmacological agents were added manually without disturbing the cover slip.

### Image quantification and statistics, software

Images were processed using LSM image examiner software (Zeiss), Image J, Fiji and by Adobe Photoshop. Quantitative analysis for Fluo4 was performed by using Fiji. Cellular size and shape were calculated manually for each image using LSM image examiner software (Zeiss). Raw data were imported into Graph pad prism for statistical analysis. Student’s t-test was done for two data set comparison. One-way ANOVA (Dunnett’s multiple comparisons test) was performed for each data set to get the significance values. To quantify the BAPTA-AM analysis we have used the Tukey’s multiple comparisons test and for Figure 6d, Sidak’s multiple comparisons test have been performed to see the significance level. Each data set was checked for the normal distribution. The P-values are as follows *** <0.001, ** <0.01, * <0.05.

### Protein expression and pull down

Expression and purification of MBP-TRPV2-Ct (C-terminal cytoplasmic domain of TRPV2 fused with MBP) and as MBP-LacZ (LacZ fused with MBP) was based on a protocol described previously^20^. The expression constructs were introduced into the *Escherichia coli* strain BL21DE3 by heat shock method. Fusion protein expression was induced by IPTG (Sigma I5502). Cells were lysed by repeated freeze-thaw cycles. The lysed extracts were cleared by centrifugation and applied to amylose resin. The resins with bound proteins were washed and the proteins were eluted with 10 mM maltose. Approximately 50 μl of amylose resin per tube with the bound MBP-TRPV2-Ct/MBP-LacZ protein was used for pull-down experiments. Soluble brain extract was added to these resins and incubated for 1 hour at room temperature (RT) in the presence or absence of Ca^2+^ (2 mM). This was followed by three washes with 200 ml PEM-S buffer. The proteins were eluted by 10 mM maltose in 100 μl solution. Eluted samples were analyzed by 10% SDS–PAGE.

### Western blot analysis

The proteins were electrophoretically transferred on PVDF membrane (Millipore IPVH00010). After blocking for 1 hour in TBST (20 mM Tris [pH 7.4], 0.9% (w/v) NaCl and 0.1% (v/v) Tween 20) containing 5% (w/v) dry skim milk, the membranes were incubated with primary antibody for 1 hours. After washing in TBST the membrane was incubated with horse radish peroxidase-conjugated secondary antibody for 1 hour at RT (25°C). Again, the membranes were extensively washed in TBST and bands were visualized on chemidoc (BioRad) by enhanced chemiluminescence according to the manufacturer’s instructions (Thermo scientific).

## Results

### Over expression of TRPV2 alters morphology of non-neuronal cells

Though the expression of TRPV2 was initially thought to be restricted to neuronal cells only, later reports confirmed the expression of TRPV2 in different non-neuronal cells also. In order to explore the effect of TRPV2 in cell morphology, we expressed TRPV2-GFP in different non-neuronal cell lines. In particular, we expressed TRPV2 in ChoKI (Chinese Hamster Ovary), HaCaT (Keratinocyte cell line), HEK (Kidney cell line), and in SaOS (primary osteosarcoma). Cells were stained with anti-tyrosinated tubulin-594 and DAPI for better estimation of the morphology of both transfected as well as non-transfected cells. In all these cases, we noted a drastic change in the cell morphology and the TRPV2 expressing cells become much flat in morphology and much bigger in sizes (**Fig 1a**). Expression of only GFP does not induce similar changes (data not shown). This indicates that TRPV2 may play important role in the regulation of sub-membranous actin cytoskeleton influencing different cellular functions such as cell adhesion and cell spreading.

**Figure 1:**
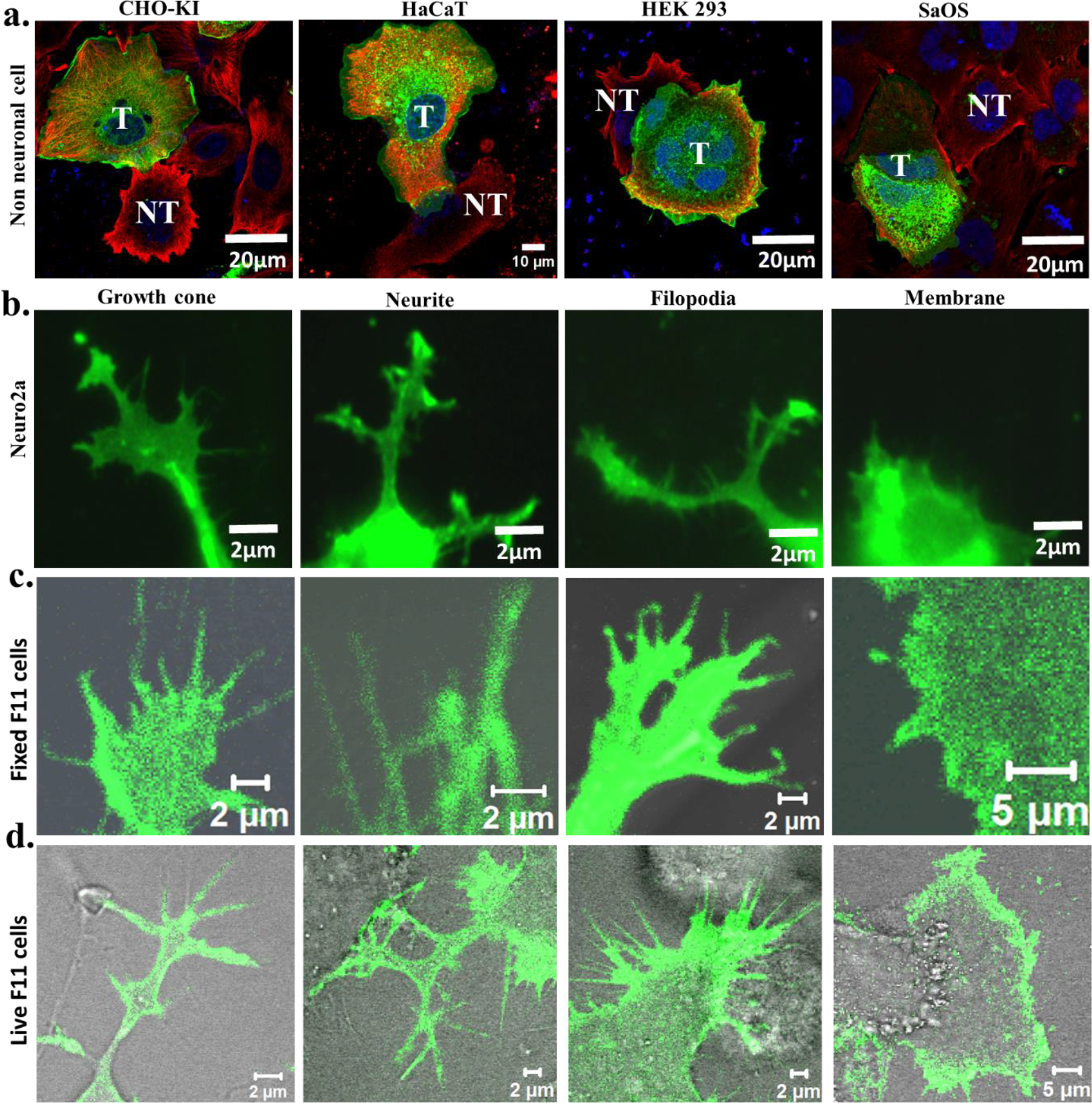
Ectopic expression of TRPV2 alters cell morphology and polarity. Shown are the confocal images of different cells or their specialized areas expressing TRPV2-GFP. **a.** Different non-neuronal cells (CHO-K1, HaCaT, HEK293, and SaOS) become enlarged after expressing TRPV2-GFP while non-transfected cells remain normal in sizes. (T and NT represents transfected and non-transfected cells respectively). All the cells were stained for Tyrosinated tubulin (Red, YL1/2 Ab) and DNA (Blue, DAPI). **b.** TRPV2-GFP localizes in the specialized cellular structures such as at growth cone, neurite and filopodia when expressed in Neuro2A (neuronal) cells. **c-d.** TRPV2-GFP localizes in the similar specialized cellular structures such as at growth cone, neurite, filopodia in fixed (c) as well as in live (d) F11 (neuronal) cells. In each case, GFP fluorescence is superimposed with the DIC images.

### TRPV2 localizes in specific membranes, neurites, growth cones and in filopodia of F11 cells

In order to understand the localization of TRPV2 in neurons, we have expressed TRPV2-GFP in Neuro2a cells, fixed after 48 hours and noted that TRPV2-GFP localizes in specific membranous regions such as in growth cones and also in filopodial structures (**Fig 1b**). Such localizations were also observed in fixed as well as in live F11 cells suggesting that these specific localizations are of special interest (**Fig 1c-d**). Growth cones are specialized structure present at the nerve endings and are involved in neurite extension, neurite bending, cell to cell contact formation and also in synapse formation^25, 35^. Indeed, TRPV2 is also present in the synaptosomes isolated from rat brain (data not shown). Filopodial structures are actin-rich membranous projections and are involved in sensing the different environment and chemical stimulus^26, 36–37^. The presence of TRPV2 in such localizations strongly suggest that TRPV2 might have importance in all these functions and may be involved in complex signaling events regulating actin dynamics and other cellular processes mediated through these structures.

### Functional TRPV2 is expressed endogenously in F11 cells

F11 cell is known to express TRPV2 endogenously^38–39^. We tried to confirm the expression of TRPV2 in F11 cells in our culture conditions. We explored the endogenous expression of TRPV2 by immunofluorescence and western blot analysis. These we performed both in presence and absence of a TRPV2-specific blocking peptide (**Fig 2a-b**). These results confirm the endogenous expression of TRPV2 in F11 cells. In order to confirm this endogenous expression further, we loaded cells with Fluo-4, a Ca^2+^-sensor dye and treated these cells with TRPV2-specific agonists and performed live cell imaging to acquire the changes in the Ca^2+^-level. Activation of TRPV2 by specific agonist (Probenecid) causes a significant increase in the Ca^2+^-level. This rise in Ca^2+^-level is transient in nature and the increased level fades off quickly. Similarly, inhibition of TRPV2 by Tranilast causes a drop in intracellular Ca^2+^-level. Further application of Probenecid causes an increase in intracellular Ca^2+^-level (**Fig 2c-d**). Quantification of multiple cells for the fluorescence intensity also confirms the same (**Fig 2e-f**). Taken together results suggest that functional TRPV2 is expressed endogenously in F11 cells.

**Figure 2:**
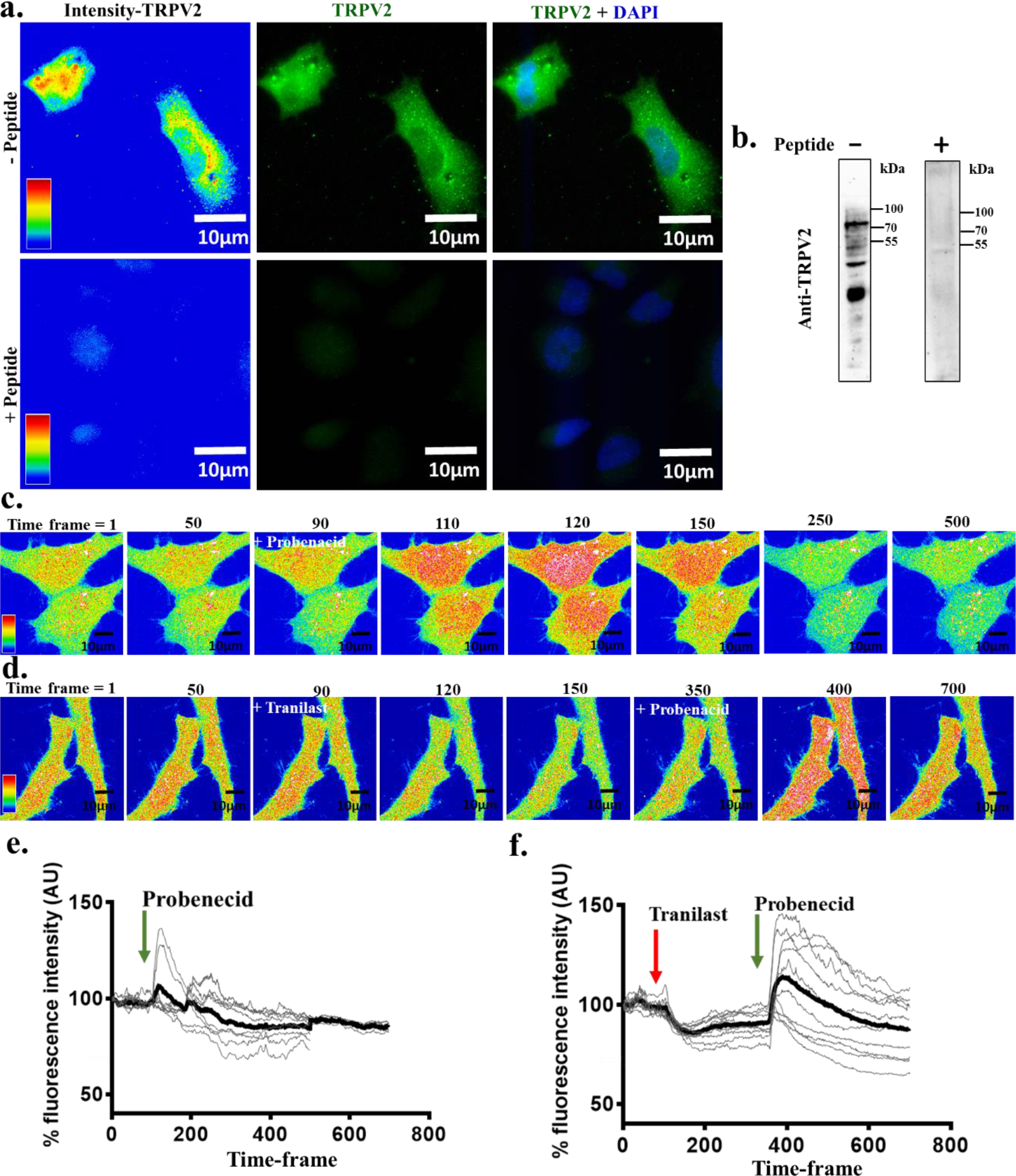
F11 cells endogenously express functional TRPV2. **a.** Immunofluorescence images of F-11 cells stained with anti-TRPV2 antibody in the absence (lower panel) or presence (upper panel) of specific blocking peptide are shown. **b.** Western blot analysis of F11 cell extract probed with anti-TRPV2 antibody are shown. The presence of specific blocking peptide diminished the TRPV2-specific immunoreactivity completely. **c.** Live cell imaging of F11 cells loaded with Fluo-4 demonstrating the transient and sharp increase in the intracellular Ca^2+^-level immediately after treating the cells with a specific activator (Probenecid, 250μM). The interval between each time frame is 0.5 sec. **d.** Similar Ca^2+^-imaging of F-11 cells shows an immediate drop in intracellular Ca^2+^-level followed by application of specific inhibitor (Tranilast, 75μM). Further application of specific activator (Probenecid, 250μM) cause again sudden increase in Ca^2+^-level. **g-h.** Quantification of intracellular Ca^2+^-levels as shown above (c-d) are represented. In each case, fluorescence intensity (in arbitrary unit) from multiple cells (n =10) are shown. The average value is shown as a thick black line. The interval between each time frame is 0.5 sec.

### Inhibition but not the activation of TRPV2-GFP results in growth cone retraction and cell retraction

The importance of TRPV1 and TRPV4 in the context of growth cone dynamics is well established^8, 22^. Therefore, we tested if TRPV2 also act in the same processes. Time-series images from TRPV2-GFP expressing live F11 cells were acquired to monitor the effect of TRPV2 activation or inhibition. For this purpose, cells that express the moderate amount of TRPV2-GFP and flatten in morphology were used. In resting conditions, these cells do not change their morphology over time drastically (within few minutes of imaging) (**Fig 3a**). However, upon inhibition of TRPV2 activity by pharmacological means results in quick retraction of cells suggesting that TRPV2 activity helps in maintenance of cell size and morphology (**Fig 3b**). Activation of TRPV2 by Probenecid results in rapid (within few seconds to minute duration) membrane ruffling, changes in lamellipodial and filopodial dynamics (**Fig 3b**). Such events often result in the initiation of neurites, branching of neurites and growth cone formation and growth cone splitting (**Fig 3d**).

**Figure 3:**
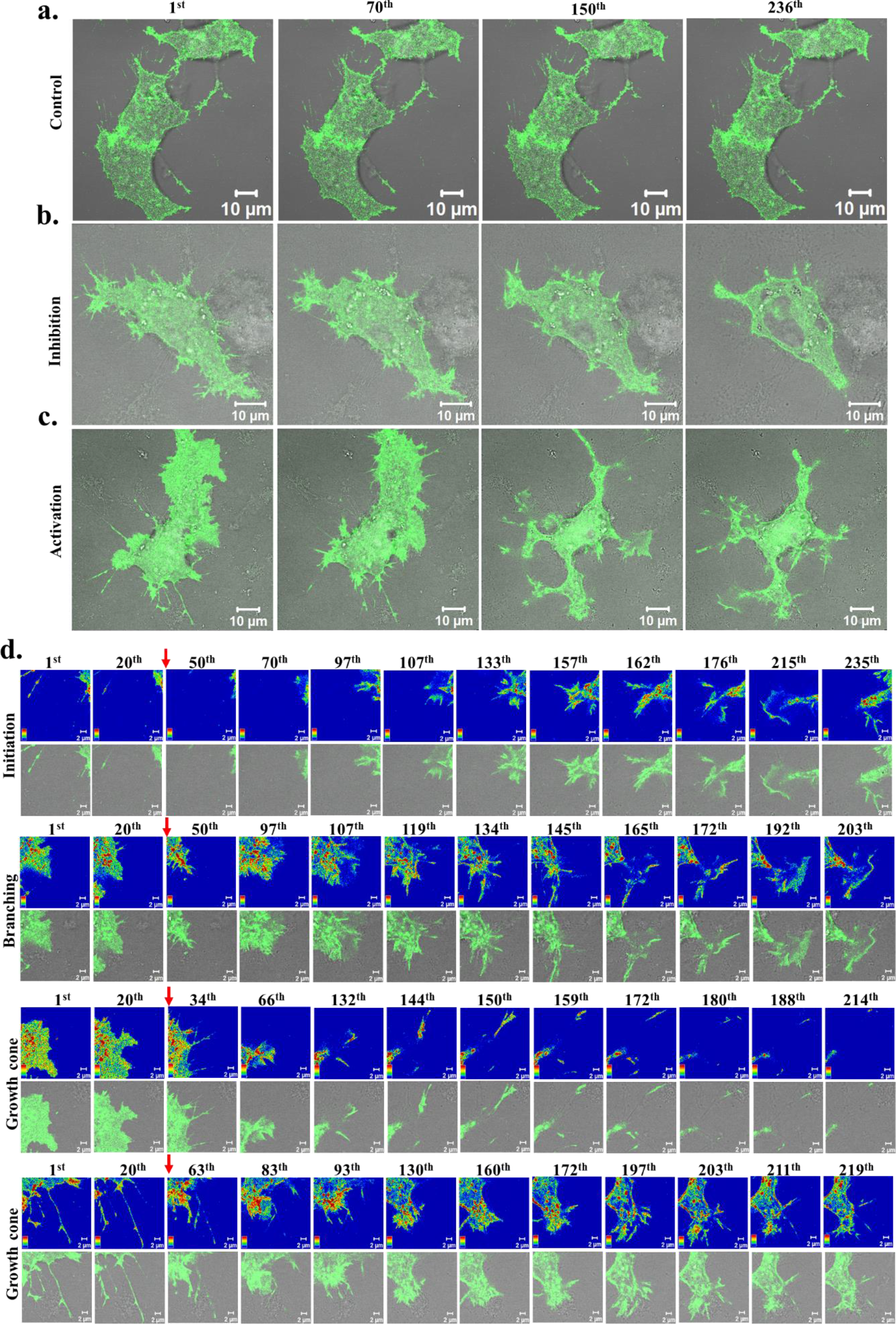
Activation of endogenous TRPV2 leads to immediate membrane ruffling and alters filopodial and growth cone dynamics. **a-c.** Representative live cell imaging of F11 cells expressing TRPV2-GFP (green) merged with DIC are shown. When cells were left un-treated (a), there is not much change in the morphology. Inhibition of TRPV2 activity by Tranilast (b) leads to a quick reduction of cell size suggesting that TRPV2 activity contributes to the maintenance of cell morphology. Activation of TRPV2 with Probenecid (c) results in rapid membrane-ruffling leading to changes in cell morphology. In each case, GFP fluorescence is superimposed with the DIC images. **d.** Shown are the time-series images of enlarged sections of a F11 cell expressing TRPV2-GFP that have been treated with Probenecid (indicated by a red arrow). Activation of TRPV2-GFP results in different events such as initiation of neurites, branching of neurites and growth cone dynamics that are controlled by sub-membranous actin cytoskeleton leading to changes in membrane ruffling. Intensity of the GFP fluorescence is represented in rainbow colors. The time gap between each image frame 0.04 Sec.

#### TRPV2 colocalizes with actin cytoskeleton and interacts with actin through its C-terminal region

As TRPV2 is present in typical structures that are enriched with actin cytoskeleton, we explored if both TRPV2 and actin colocalizes. For that purpose, we immunostained TRPV2 and labelled actin cytoskeleton by Phalloidin. We noted that TRPV2 co-localizes with actin cytoskeleton in the specific regions such as in filopodia and growth cone regions (**Fig 4a**). To explore further if TRPV2 co-localizes with actin cytoskeleton in live cell, we co-expressed TRPV2-GFP and actin-RFP and performed live cell imaging. TRPV2-GFP and Actin-RFP colocalizes in all these specific cellular areas (**Fig 4b)**.

**Figure 4:**
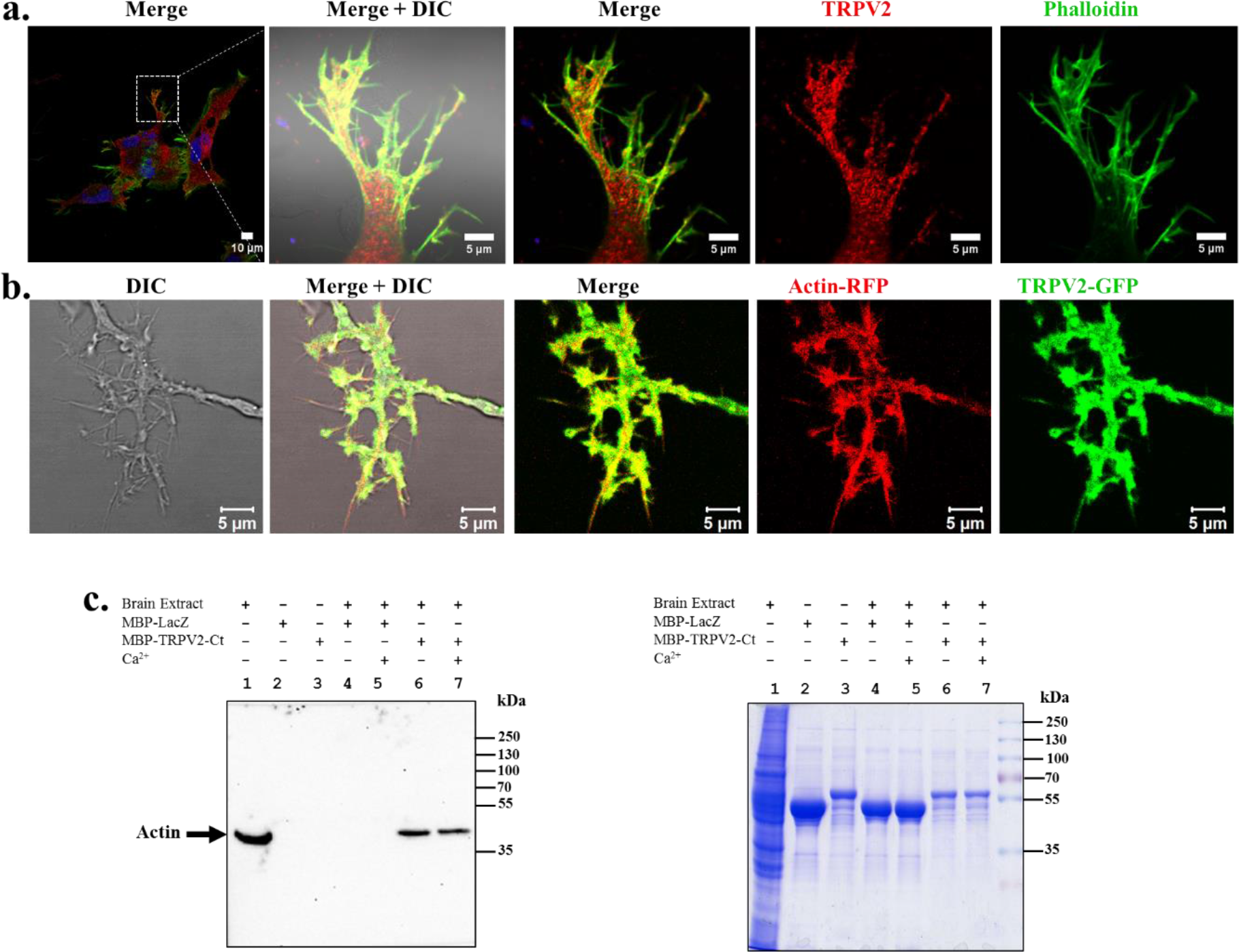
TRPV2 colocalizes with sub-membranous actin cytoskeleton and interacts with actin. **a.** TRPV2 co-localizes with Phalloidin in filopodia and specific cellular areas. **b.** Live cell image of a F11 cell co-expressing TRPV2-GFP (green) and actin-RFP (red) demonstrating colocalization of TRPV2-GFP and actin-RFP in specific cellular regions such as in neurites and in filopodia. **c.** The C-terminus of TRPV2 interacts with actin. Soluble brain extract supernatant (lane 1) was pulled down with purified MBP-LacZ (lane 4-5), MBP-TRPV2-Ct (lane 6-7) and probed for bound actin by Western blot analysis.

Next, we tested if TRPV2 interacts with actin. For that purpose, we expressed the C-terminus of TRPV2 as a MBP-tagged protein and performed a pull-down experiment with cattle brain extract. The pull-down samples were probed for the presence of actin by western blot analysis. Actin-interacts with MBP-TRPV2-Ct but not with MBP-LacZ, both in presence and absence of Ca^2+^ (**Fig 4c**). This confirms that the C-terminus of TRPV2 interacts with actin.

#### Activation of TRPV2 causes membrane remodeling

Next, we explored if TRPV2 activation and inhibition can cause rapid changes in the cell morphology and if such changes accompany remodeling of sub-membranous actin cytoskeleton. Activation of TRPV2 by Probenecid leads to rapid changes in actin cytoskeleton causing changes in cell morphology, such as extension of cell membrane, formation of massive lamellipodium and often merging of lamellipodium (**Fig 5a**). Such effects can also be reproduced by application of 2APB (10μM), another activator of TRPV2 (**Fig 5c**). In all cases, such changes are accompanied by rapid translocation of TRPV2 in the membrane, alteration in the actin filaments. Activation of TRPV2 induce rapid translocation in leading edges but not on the filopodial tips (**Fig 5c**). These results strongly suggest that TRPV2 can be functional for cellular events, such as different aspects of neuritogenesis that involves actin cytoskeleton.

**Figure 5:**
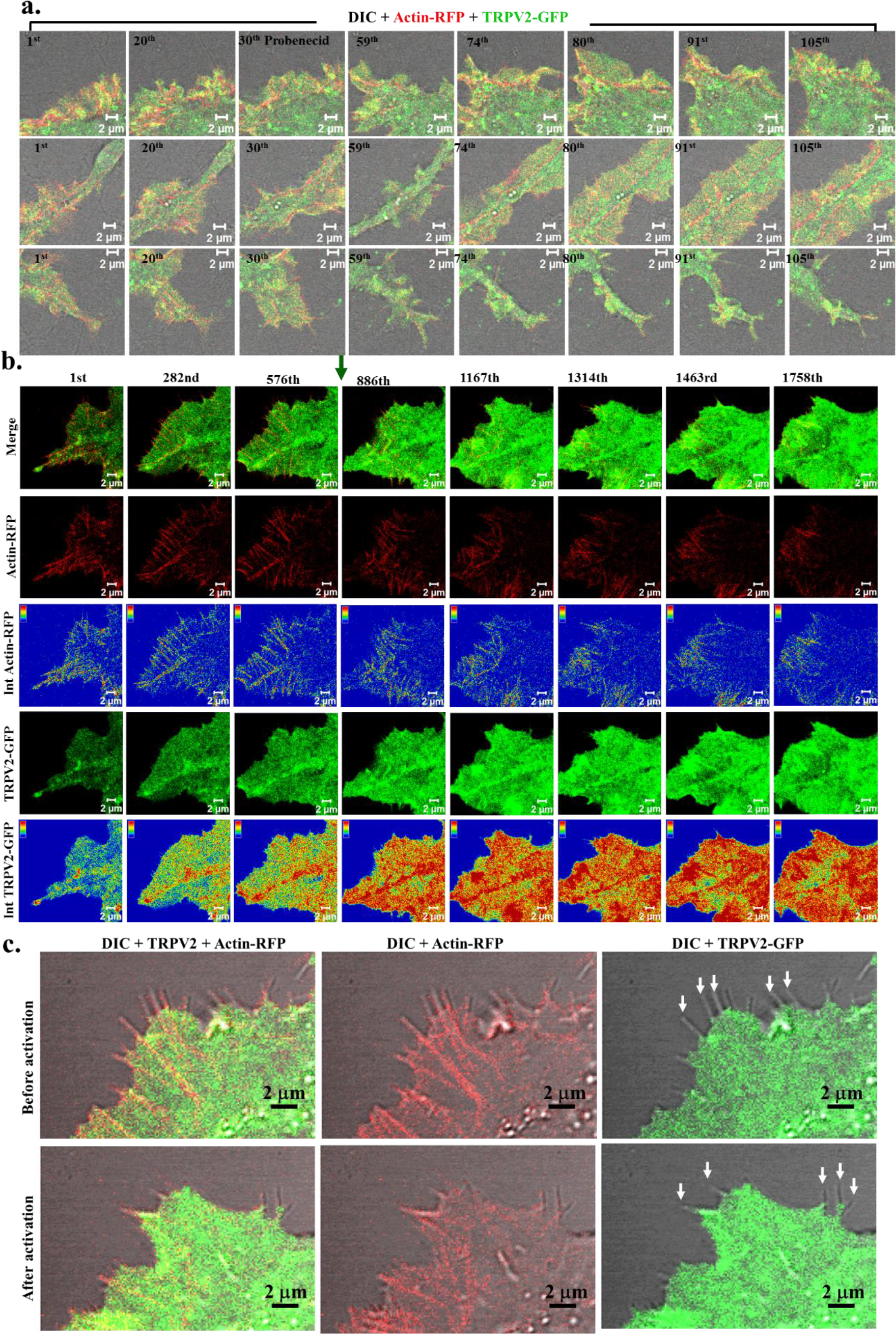
Activation of TRPV2 induce rapid translocation in leading edges but not on the filopodial tips. **a.** Live cell image of F-11 cell co transfected with TRPV2-GFP and actin-RFP shows that activation of TRPV2 by Probenecid alters actin cytoskeleton dynamics leading to changes in cell morphology, such as extension of cell membrane (upper panel), formation of massive lamellipodium (middle panel) and merging of lamellipodium (lower panel). **b.** Activation of TRPV2-GFP by 2APB (indicated by green arrow at 886^th^ frame) also causes changes in sub-membranous actin cytoskeleton and results in rapid translocation of TRPV2-GFP to the leading edges, merging of actin-ribs in the lamellipodium boundary. Intensity profile of the TRPV2-GFP and actin-RFP are provided in below. **c.** Shown are the enlarged portion of the leading edge of a F11 cell expressing TRPV2-GFP and actin-RFP before and after activation with 2APB. The filopodial tips are marked with white arrows. TRPV2-GFP is mainly present in the filopodial base but not in the filopodial tips.

### Exogenous expression of TRPV2 cause changes in cell morphology and induces neuritogenesis

To explore the importance of TRPV2 in functions related to neurite formation and further extension, we transfected TRPV2-GFP in neuronal cells and analyzed the different properties of neurites originated from non-transfected cells as well as cells transiently expressing TRPV2-GFP. Neuro2a cells expressing TRPV2-GFP become much elongated and long neurites are visible (**Fig 6a**). F11 cells overexpressing TRPV2-GFP become much elongated compared to non-transfected cells and drastic difference in the cell length is visible (**Fig 6b**). To explore if TRPV2 overexpression can indeed enhance neuritogenesis, we quantified the percentage of cells that show the presence of at least one single primary (1°) neurite, or at least two primary neurites (both originated from the cell body) or even more number of primary neurites (all originated from the cell body). Approximately 70% of TRPV2-GFP expressing F11 cells develop at least one single neurite within 24 hours. In contrast, only ~25% non-transfected F11 cells develop at least one neurite within the same time points. Similarly, ~88% cells expressing GFP-only have no primary neurites. The percentage of cells with at least two or more than two primary neurites are much higher in cells that are expressing TRPV2-GFP than that of the non-transfected F11 cells or GFP-only expressing cells (**Fig 6c**).

**Figure 6:**
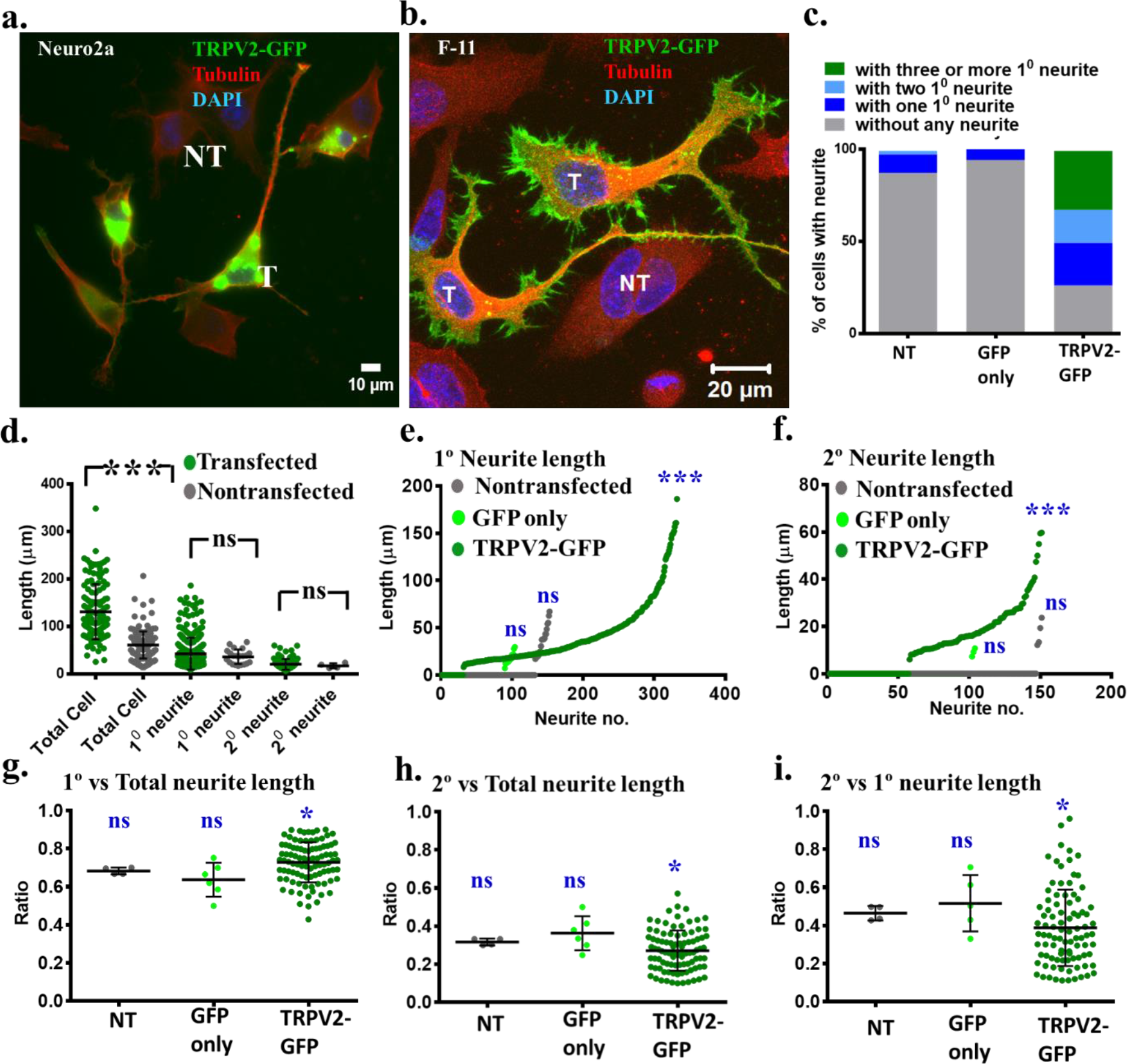
Exogenous expression of TRPV2 induces neuritogenesis and enhances cell elongation. **a-b.** Representative fluorescence microscopic images demonstrating Neuro2A (a) and F11 (b) cells expressing TRPV2-GFP. Transfected cells (T) become much elongated and have a higher number of neurites with complex branches compared to non-transfected (NT) cells. **c.** Expression of TRPV2-GFP enhances neuritogenesis. Percentage of F-11 cells having at-least one primary neurite, two primary neurites, more than 2 primary neurites or no neurites were quantified (n =149 for non-transfected cells, 104 for GFP expressing cells and n =131 for TRPV2-GFP expressing cells). **d.** Quantification of the length of the entire cell (n = 149), primary (1°) neurite (n = 100) and secondary (2°) neurites (n = 80) originated from F11 cells over expressing TRPV2-GFP are shown. The mean cell length becomes significantly different (p-value) while the mean length of the 1° and 2° neurites are not-significantly different. **e-f.** Length of the 1° neurites and 2° neurites (n =131 TRPV2-GFP expressing cells, 104 GFP expressing cells, and n = 149 non-transfected cells) present in F11 cells are plotted in ascending orders. For cells without any 1° neurite, the length of the 1° neurite is considered as zero (e). Similarly, for 1° neurites without any 2° neurite, the length of the 2° neurite is considered as zero (f). **g-i.** Ratio (of length) of 1° neurites to total cell (g), 2° neurite to total cell (h) and 1° to 2° neurite (i) are plotted for each cell. Marginal differences in these ratios between TRPV2-GFP expressing cells and GFP only expressing cells or non-transfected cells are observed. P values ≤ 0.001, 0.01, 0.5, 0.1 are considered as ***, **, *, and ns respectively.

Subsequently, we analyzed for the length of the primary and secondary neurites. TRPV2-GFP expressing cells have more number of primary (1°) as well as secondary (2°) neurites (**Fig 6d-f**). However, we noted that the average length of the 1° or 2° neurites are almost same (and the difference are non-significant) between cells expressing TRPV2-GFP or that are non-transfected (**Fig 6d**). Further analysis reveals that the length of the 1° neurites consists majority of the length of the total neurites and the value is marginally more for TRPV2-GFP expressing cells (average value is ~70%) than non-transfected cells (**Fig 6g**). The ratio of 2° vs. total neurite length also suggests that 2° neurites contribute ~25% of the total neurites length and such value is marginally less for TRPV2-GFP expressing cells than that of the non-transfected cells (**Fig 6h**). Similarly, the ratio of 2° vs. 1° neurites also indicates that the average value is slightly less for TRPV2-GFP expressing cells than that of the non-transfected cells (**Fig 6i**). In all these cases, expression of only GFP does not induce neurites and these cells are comparable to non-transfected cells only (**Fig 7a**).

**Figure 7.**
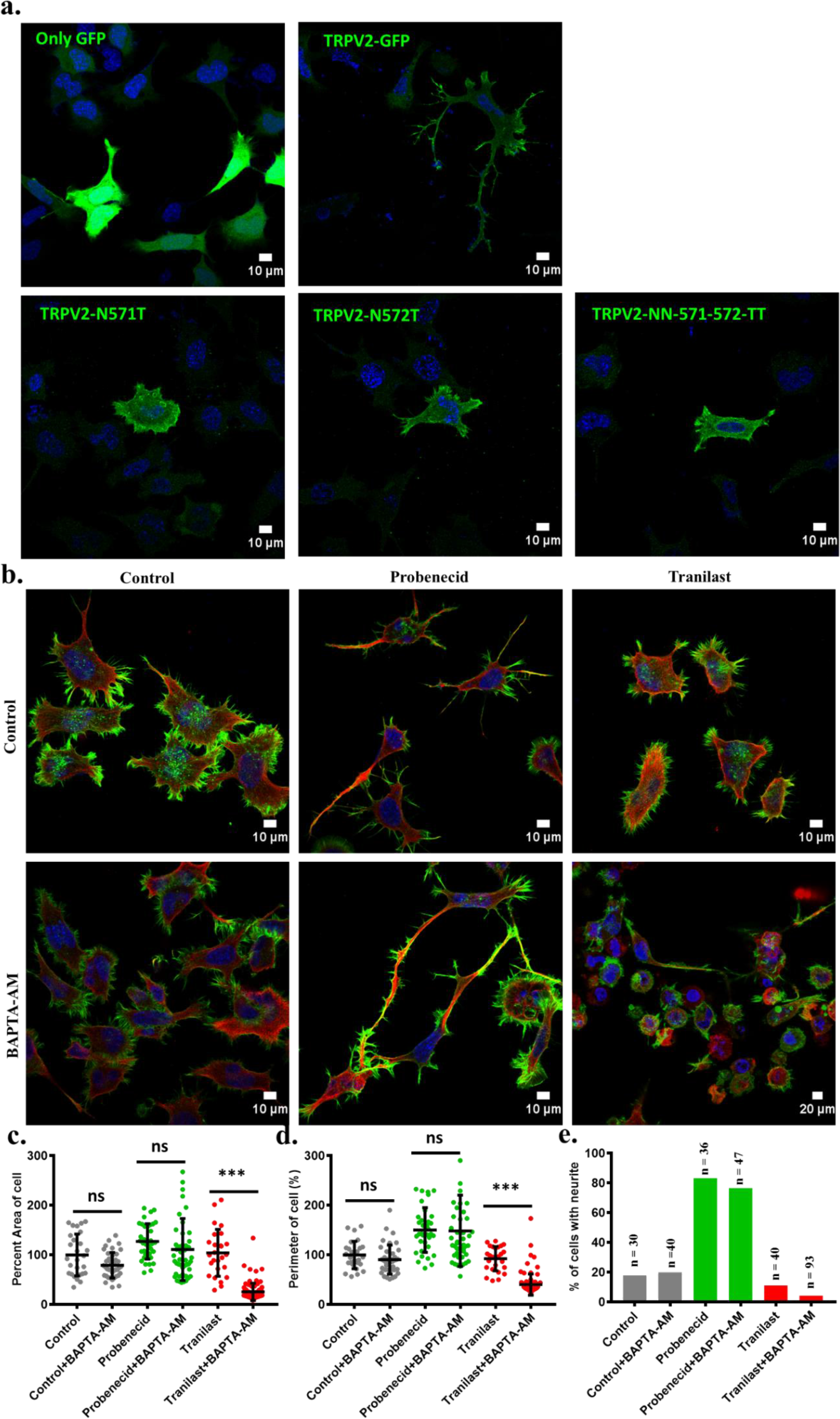
TRPV2-mediated neuritogenesis is dependent on wild type TRPV2 but independent of intracellular Ca^2+^-levels. **a.** Only Wild-type TRPV2-GFP induces long neurites. Shown are the F11 cells expressing TRPV2-Wt-GFP, or TRPV2-mutants or GFP only. Cells were stained with DAPI and images were acquired by confocal microscopy. **b.** F11 cells were grown in absence or in presence of BAPTA-AM and were either treated with TRPV2 activator (Probenecid) or TRPV2 inhibitor (Tranilast). Cells were stained with Phalloidin (Green), Microtubules (Red) and DAPI (Blue).

This data strongly suggests that over-expression of TRPV2-GFP induces more number of neurites per cell (and mainly from the cell body), but does not affect average length of the individual neurites *per se*. Alternatively, this data suggest that over-expression of TRPV2 induces signaling events that cause initiation of more neurites from the cell body, but once the neurites are formed, their lengths are independent of the level of TRPV2 expression *per se* (discussed later).

### TRPV2-mediated neuritogenesis is dependent on wild-type TRPV2 and is independent of intracellular Ca^2+^-levels

In order to understand the importance of TRPV2 in the neuritogenesis, we expressed wild type as well as TRPV2 mutants (TRPV2-N571-T, TRPV2-N572-T, TRPV2-NN571-572-TT) which are defective in their properties^33^. We observed that wild type TRPV2 but not the mutants induce long neurites (**Fig 7a**). We explored if endogenous TRPV2 also involved in the neuritogenesis and if pharmacological activation or inhibition of TRPV2 alters this. We also tested if such neuritogenesis is dependent on the intracellular Ca^2+^ levels. For that we have used F11 cell in presence or absence of BAPTA-AM and modulated TRPV2 by activator or inhibitor. We noted that F11 cells induce neurites due to TRPV2 activation even in the presence of BAPTA-AM (**Fig 7b**). In fact, presence of BAPTA-AM does not make any differences in different cellular properties, such as percent area of cell, percent perimeter of cell or even percentage of cells with neurites (**Fig 7c**). However, cell spreading is significantly reduced when TRPV2 is inhibited in presence of BAPTA-AM. These results suggest that TRPV2-induced neuritogenesis involves signaling events that are dependent on wild type TRPV2 but independent of intracellular Ca^2+^.

### Endogenous TRPV2 in F11 cells is involved in neuritogenesis

In order to test the importance of TRPV2 in neuritogenesis events in details, we have activated endogenous TRPV2 with a specific activator (Probenecid) at a sub-optimal concentration (10μM) for 12 hours. Similarly, endogenous TRPV2 activity was also inhibited by using Tranilast at 75μM concentration (more than optimal concentration to ensure the complete blockade of TRPV2 activity) for 12 hours. Subsequently, cells were fixed without disturbing the media and the cells were immunostained with tubulin antibody to analyze their morphology and length of the neurites as well. Our results suggest that activation of TRPV2 leads to elongation of cell sizes very much whereas pharmacological inhibition of TRPV2 reduces the cell length significantly compared to the control cells. We also measured several parameters related to neuritogenesis events such as the percentage of cells having no-neurite, or percentage of cells having at least one 1° neurite or more than one 1° neurites (originated from the cell body). We also measured other parameters such as total cell length, the length of primary neurites and length of secondary neurites and their comparative ratios.

In control condition, there are around 30% cells which are without any neurite while TRPV2-activated (Probenecid-treated) conditions only 5-7% of cells are without any neurite (**Fig 8a**). In case of TRPV2 inhibition (Tranilast-treated cells), around 60% cells remain without any neurite, suggesting that activation of endogenous TRPV2 enhances the neuritogenesis process as a whole (enhances the % of cells with neurites) while inhibition of TRPV2 reduces the % of cells with neurites). In a similar manner, the percentage of cells with a single primary neurite is increased when TRPV2 is activated (35%) compared to cells that were grown in resting conditions (20%). Similarly, the percentage of cells with at least one secondary neurite (originated from primary neurite) is increased after activation of endogenous TRPV2 (55%) compared to cells that are in resting condition (40%) or grown in inhibited conditions (3-4%). This result also strongly establishes a direct correlation between TRPV2 function with the initiation of neurites from cell body or even from the primary neurite. In addition, the length of the primary neurites is significantly longer in cells that were treated with TRPV2 activator and significantly shorter in cells that are treated with a TRPV2 inhibitor (**Fig 8b** and **Fig 8e**). However, analysis of the length of secondary neurites indicates that there is no significant difference between cells that are pharmacologically modulated (activated or inhibited) with the cells that are grown in control conditions (**Fig 8c** and **Fig 8f**). These results may suggest that endogenous TRPV2 activity is involved in the formation of cell polarity, at least in the F11 cell. This result suggests that though activation of endogenous TRPV2 induces more secondary neurites, their lengths are independent of TRPV2 function and therefore suggest the involvement of TRPV2 in complex signaling events.

**Figure 8:**
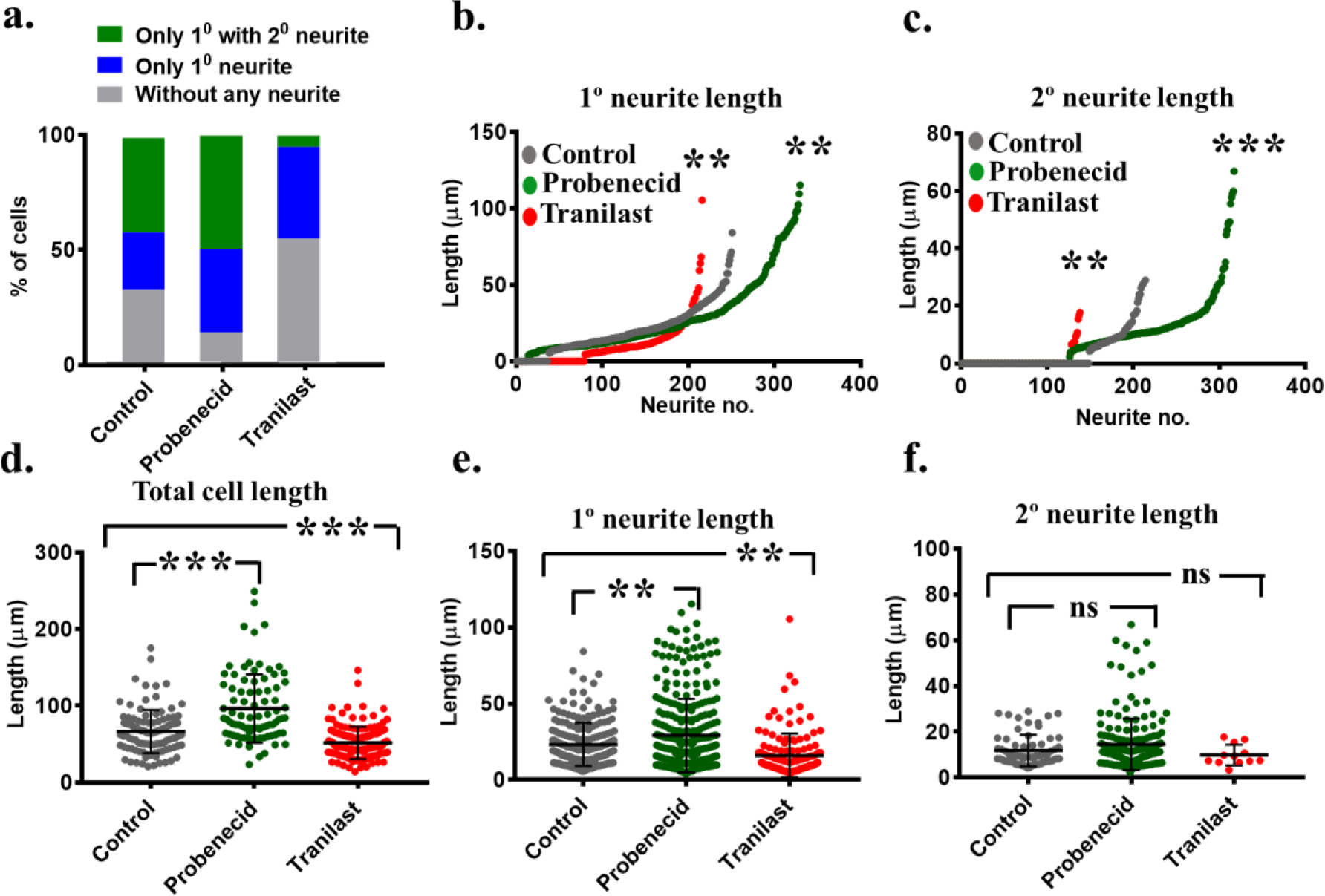
Endogenous TRPV2 activation increases cell size and enhances neuritogenesis. **a.** Activation of TRPV2 induces more neurites. The F-11 cell was grown in control condition, activated with Probenecid (10μM for 12 hours) or inhibited with Tranilast (75μM for 12 hours). Cells were quantified and percentage of cells without any neurite, with at-least one primary (1°) neurite, and with at-least one secondary (2°) neurite originated from primary neurite are plotted. **b.** Length of the 1° neurites (n =109 for control, 89 for Probenecid-treated conditions and 143 for Tranilast-treated conditions are plotted in ascending orders. For cells without any 1° neurite, the length of the 1° neurite is considered as zero. **c.** Length of the 2° neurites (n =109 for control, 89 for Probenecid-treated conditions and 143 for Tranilast-treated conditions) are plotted in ascending orders. For 1° neurite without any 2° neurite, the length of the 2° neurite is considered as zero. **d-f.** Length of the total cell (d), 1° neurite (e) and 2° neurite (f) from cells grown in control condition, activated with Probenecid or inhibited with Tranilast are plotted. The average length of the 2° neurites is non-significantly (ns) different when cells were grown in different conditions. P values ≤ 0.001, 0.01, 0.5, 0.1 are considered as ***, **, *, and ns respectively.

To establish the importance of TRPV2 in neuritogenesis further, we analyzed the ratio between primary, secondary and total neurite lengths. We noted that the ratio of primary neurite length with total neurite length is significantly higher when cells were treated with TRPV2 activator and remain unchanged when cells were treated with inhibitors (**Fig 9a**). In the same notion, the ratio of secondary neurite length with total neurites length is significantly lower when cells were treated with TRPV2 activator and remain unchanged when cells were treated with inhibitors (**Fig 9b**). A similar trend in the ratio between the lengths of secondary neurite with primary neurite is observed (**Fig 9c**). This result suggests that endogenous TRPV2 affects primary neurites dominantly than the secondary neurites in terms of length.

**Figure 9:**
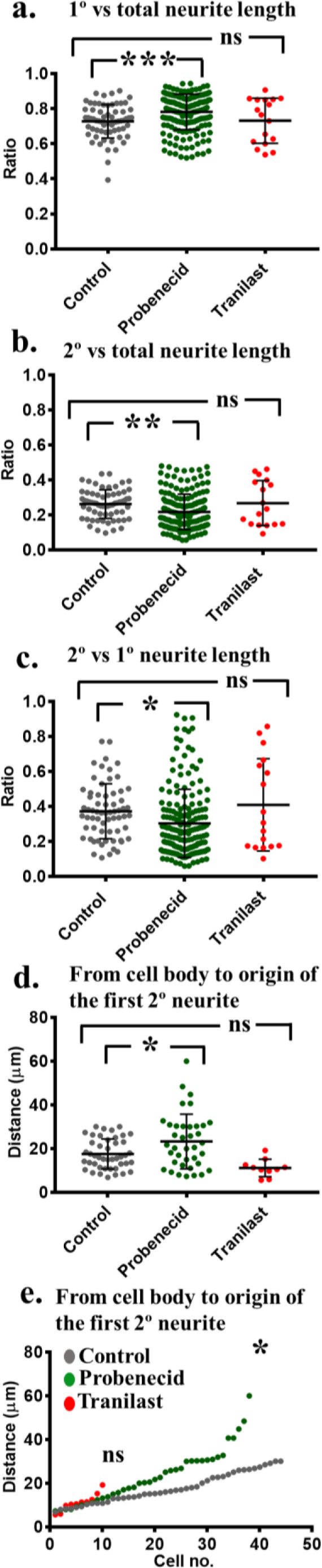
Activation of endogenous TRPV2 affects primary neurites more than secondary neurites. **a-c.** The ratio of primary neurite length to total cell length (a), or secondary neurite length to total cell length (b) or secondary neurite to primary neurite length were quantified for cells grown in different conditions. This ratio is significantly higher when cells are grown in presence of TRPV2 activator Probenecid (10μM for 12 hours) and mean is unaltered when grown in presence of inhibitor Tranilast 75μM for 12 hours (n =109 for control, 89 for Probenecid-treated conditions and n = 143 for Tranilast-treated conditions). **d-e.** Cells were grown in different conditions and the distance between cell body to the origin of first secondary neurite developed was quantified. Activation of TRPV2 increases the distance (of primary neurite) between cell body to the first point where secondary neurite originates. The mean/average distance is unaltered when grown in presence of inhibitor Tranilast. The values were plotted in ascending order (e). The P values ≤ 0.001, 0.01, 0.5, 0.1 are considered as ***, **, *, and ns respectively.

### TRPV2 affects neurite branching

The changes in number, as well as the length of primary and secondary neurites in general, suggest that TRPV2 is involved in the initiation of new neurites. Therefore, we tried to explore the importance of TRPV2 in neurite branching process. In order to explore this, we have measured the distance between the cell body and the origin of the first secondary neurite (**Fig 9d-e**). Such analysis indicates that activation of TRPV2 significantly increases the primary neurite length between the cell body and the origin of the first secondary neurite. In the same notion, inhibition of TRPV2 also reduces this distance. This result establishes a correlation between TRPV2 activities with the initiation of new neurites.

### Activation of TRPV2 results in increased cAMP level in neurites and in neurite branching points

Changes in the Ca^2+^ concentration and cAMP levels in the growing and /or undifferentiated neurite primarily determines which growing neurite will form the axon^1,7^. We tested if TRPV2 modulation enhances the neurite outgrowth and branching by altering cAMP signaling. Therefore we tested the level of CREB and phospho-CREB levels in control condition as well as long term TRPV2 activated or inhibited conditions. The total level of CREB remain unaltered and increased in case of inhibition as analyzed by western blot analysis (**Fig 10a**) or by FACS-based analysis (**Fig 10b**). However, the total level of phospho-CREB increases in case of activation and much more in case of inhibition as analyzed by western blot analysis (**Fig 10a**) as well as by FACS-based analysis (**Fig 10c**).

To analyze if TRPV2 activation can alter cAMP levels as a quick response, we have used a FRET-based cAMP-sensor and performed live cell imaging experiments (**Fig 10d**). This sensor shows loss-of-FRET signal upon cAMP interaction. Therefore “loss-of-FRET” signal indicates for increased cAMP levels (**Fig 10d-g)**. During live cell imaging after TRPV2 activation with Probenecid, we found a drastic decrease in the FRET signal suggesting that TRPV2 activation results in increased cAMP concentration in the F11 cell (**Fig 10f**). Such increased cAMP level is also observed in the specific regions of neurites which later on split and become a branching point (**Fig 10g**).

**Figure 10:**
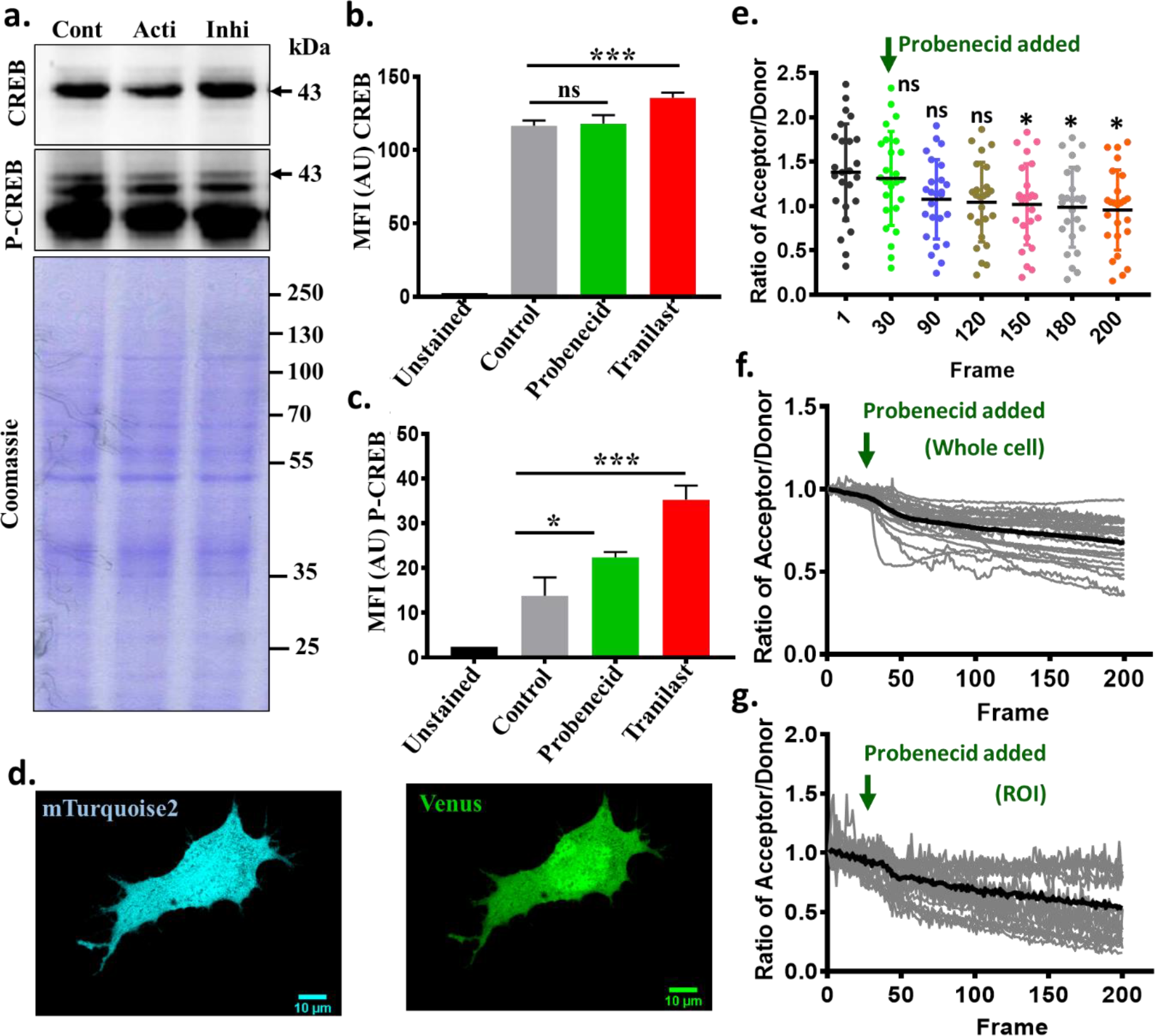
Activation of endogenous TRPV2 increases cAMP level in F-11 cells. **a.** Western blot analysis of F11 cell extracts for CREB and phospho-CREB levels. F11 cells were either grown on control conditions or treated with activators or inhibitors for 12 hours before the samples were prepared. **b-c.** F11 cells grown in control conditions or treated with activators or inhibitors for 12 hours. Cells were analyzed for CREB and phospho-CREB levels by FACS analysis. Activation of TRPV2 does not alter CREB levels much, but alters phosphor-CREB levels (n = 3). **d.** F11 cells were transfected with FRET-based cAMP-sensor molecule (Epac-SH188) and the respective confocal images acquired in different spectra are shown. **e.** Multiple F11 cells transfected with the cAMP sensor were quantified and the ratio of acceptor/donor values are shown. Each dot represents the “absolute values” obtained from a single cell. Live F11 cells were imaged for total 200 frames (≈ 5 Min in total, time interval between each frame is 0.5 sec) and cells were activated with Probenecid at 25^th^ Frame. Decreased values suggest increased cAMP formation after addition of TRPV2 activators. P values that are ≥ 0.1 and ≤ 0.01 are considered as non-significant (ns) and * respectively. **f-g.** Shown is the ratio of acceptor/donor in live F11 cell transfected with the cAMP sensor. Cells were activated with Probenecid at 25^th^ Frame. The initial value of the ratio is normalized and considered at 1. Gradual decline in values suggest increased cAMP formation after addition of TRPV2 activators, both in whole cell (f) and in specific ROI drawn on the neurites and branching points (g).

Taken together our results suggest that TRPV2 interacts with actin and activation of TRPV2 results in reorganization of actin cytoskeleton, altered cAMP levels which in turn trigger enhanced neuritogenesis.

## Discussion

Regulation of sub-cellular cytoskeleton and vesicle recycling are essential for several cellular functions, such as cell adhesion, cell spreading, cell-cell contact formation, and proper cell signaling events^40–45^. In the case of neuronal cell, specific functions such as filopodial dynamics, growth cone formation, neurite initiation, neurite extension, neurite turning, and neurite branching are critical, and relevant for neuronal plasticity^46–47^. Such processes are important for neuron-neuron contact formation and precise neuronal circuit formation which is essential for several physiological and sensory functions^47^. Indeed, such processes are very precise and regulated by multiple factors and often mis-regulation of such processes leads to the development of different yet common pathophysiological conditions. For example, insufficient neuritogenesis may trigger neurodegeneration while excess sprouting may lead to hypersensitivity^48–49^.

Different ion channels, such as K^+^ and Na^+^ channels are involved in the neuritogenesis events^50–51^. Never-the-less, different steps of neuritogenesis are largely dependent on Ca^2+^-signaling events and thereby involved different Ca^2+^ channels^2–3,7^. A large number of reports confirmed that different physical and chemical cues affect the process of neuritogenesis significantly^11^. As TRPV members are Ca^2+^-permeable channels, polymodal in nature, regulated by different physical as well as chemical cues, external factors and also by endogenous factors, TRPV channels are ideal candidates that can detect different chemical cues, integrate different signals and therefore are suitable for functions related to neuritogenesis (**Fig 11**). Indeed, previous studies have shown that many of these signaling events are Ca^2+^ dependent as well as Ca^2+^ independent. In our experimental conditions we found that endogenous TRPV2 modulation in presence or absence of BAPTA-AM results in the morphological change. Therefore our data strongly suggest that TRPV2 in involved in both Ca^2+^-dependent as well as Ca^2+^-independent signaling events.

**Figure 11:**
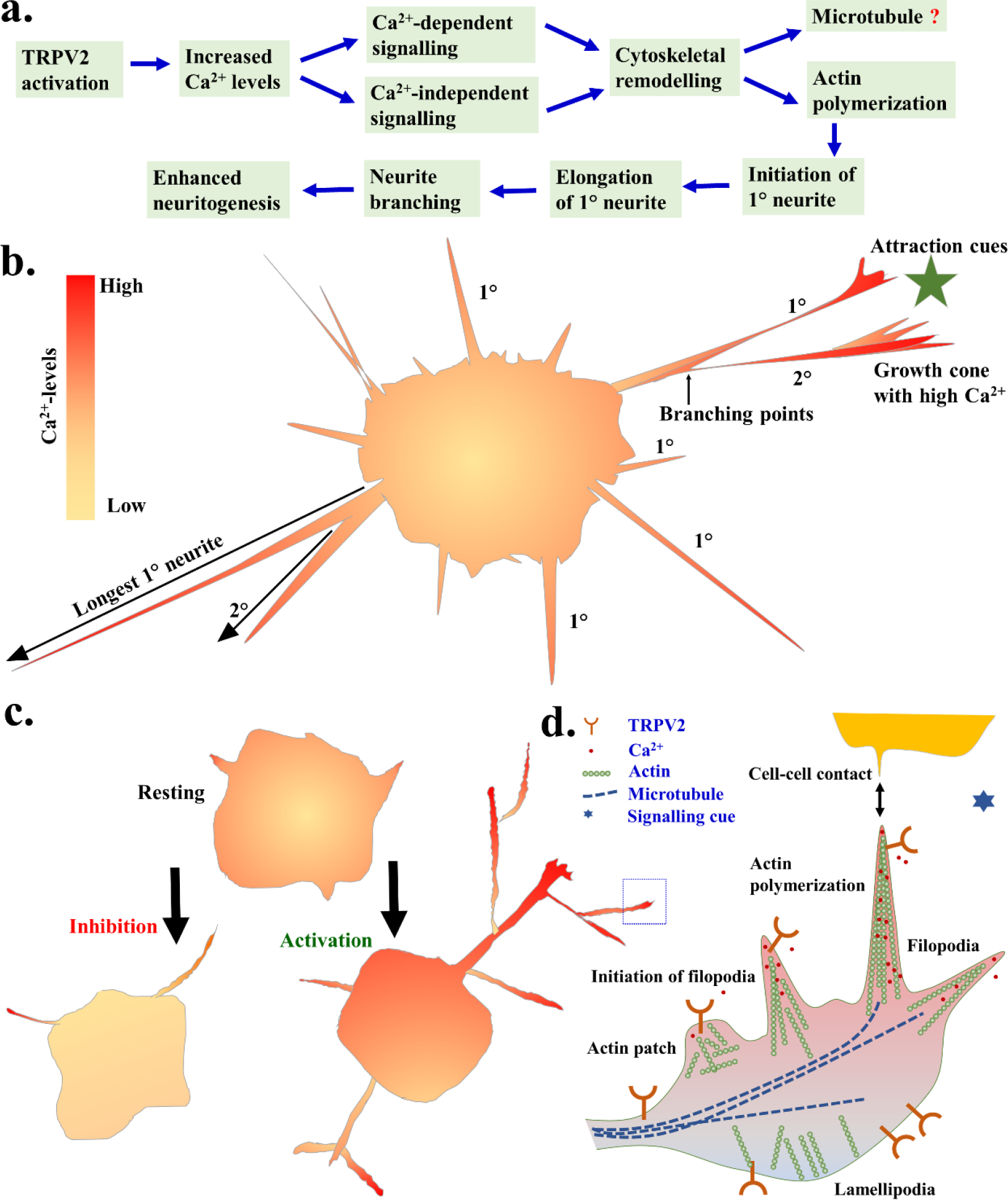
Schematic representation of plausible cellular and molecular events regulated by TRPV2 leading to enhanced neuritogenesis. **a.** TRPV2 activation induces a cascade of cellular events involving both Ca^2+^-dependent and Ca^2+^-independent signaling leading to cytoskeletal remodeling. TRPV2 activation induces new 1° neurites, branching events and induce new 2° neurites leading to enhanced neuritogenesis. However, once initiated, further extension of the formed neurites seem to be independent of TRPV2 activation. **b.** Schematic representation of a neuronal cells with neurites and their relative intracellular Ca^2+^-levels. TRPV2 seem to contribute to the regulation of Ca^2+^-homeostasis in different parts of the cells and especially in the leading ends where Ca^2+^-levels are usually high. **c.** Shown is a simplified model demonstrating the changes in cell morphology and development of complex neurites due to activation of TRPV2. **d.** A simplified model represents the involvement of different molecular factors such as TRPV2, actin, microtubule and Ca^2+^ in specialized subcellular structures (indicated by dotted box) such as in filopodia and in neuronal growth cone.

In this context, it is important to mention that cAMP also plays a key role in the neuritogenesis. Level of cAMP in the growing neurite and cAMP signaling through positive feedback mechanism determines the fate of undifferentiated neurite and neurite branching^1,7.^. This is in agreement with our results here activation of TRPV2 results in the increased cAMP concentration and neurite branching. The increment in phosphor-CREB level due to TRPV2 activation is observed in long term activation. However, our FRET-sensor based experiments detect a more direct increment of cAMP levels due to TRPV2 activation and such changes can be observed in very short term, such as in minutes after TRPV2 activation.

Expression of TRPV2 in sensory and motor neurons has been reported in the early stages (E10) of neuronal development^52^. Previously, expression of TRPV2 in specific regions of the brain and in sensory neuron have also been reported^53–54^. In such regions, TRPV2 mainly localizes in neurites and in nerve ends where TRPV2 colocalizes with different synaptic vesicular markers^55^. TRPV2 is also involved in the neuronal contact formation within non-neuronal tissues. For example, of the small CGRP^+ve^ neurons projecting to the skin, 11.6% were positive for TRPV2. Involvement of TRPV2 in neuro-protection have also been reported^56^. Here we demonstrate that TRPV2 plays important role in the regulation of neuritogenesis process. Our results are in line with previous reports which demonstrate the involvement of other members, namely TRPV1 and TRPV4 in similar functions^8–9, 21–23, 57^.

Our work also accords well with previous works demonstrating the importance of TRPV2 in neuritogenesis in general^10, 27^. However, these previous studies are primarily done in PC12 cells, performed in presence of certain growth factors (such as NGF or BDNF), and often TRPV2 was activated by mechanical stimuli or by membrane stretching^10, 27^. As these growth factors and/or mechanical stimuli can activate a plethora of signaling events, from previous experiments it is difficult to derive the actual involvement of TRPV2 in such neuritogenesis process. Previously it has been reported that knockdown of TRPV2 by siRNA results in a reduction in NGF-induced neurite outgrowth, at least in *in vitro* culture conditions^10^. However, TRPV2 knockout animals have normal sensory functions suggesting that the neuronal circuits are not altered by the deletion of TRPV2 gene *in vivo*^58^. These facts may also suggest that importance of TRPV2 in neuritogenesis is not essential, and TRPV2 functions are pleiotropic in nature. Never-the-less, the experiments we described in this work are from *in vitro* cell cultures mimicking peripheral neurons and are free from other added growth factors. All the experiments were mainly performed within undisturbed media conditions. Thus, the results described in this work represent the involvement of TRPV2 in neuritogenesis process, specifically in response to pharmacological modulation only and in the absence of added growth factors or mechanical stimulation-induced activation of TRPV2 (as well as other receptors/channels). Our results establish that activation of TRPV2 induces neurite initiation and branching. However, once the neurites are initiated, the actual lengths are independent of the TRPV2 expression level. In this context, it is important to mention that involvement of TRPV2 in the neurite initiation and neurite branching seem to be a complex process that can be enhanced by TRPV2 activation in general, but certain parameters are unchanged by inhibition of TRPV2. Such process may involve other signaling pathways as well.

Filopodial tips response to mechano-chemical stimuli, micro-heating, suggesting that chemical, temperature and mechano-sensing protein complexes are present there^13, 59–61.^ Considering the fact that TRPV2 can be activated by temperature, mechanical stimuli and other chemical factors, our observation that TRPV2 is localized in the filopodial structures is in full agreement with the different sensory properties that filopodial structure possesses. Previously we have reported the presence of other TRPV members, such as TRPV1 and TRPV4 in the filopodial structures too. However, the localization and function of TRPV2 in these structures are not exactly similar to TRPV1 or TRPV4. For example, TRPV1 activation results in retraction of growth cone while activation of TRPV2 does not induce the growth cone retraction. In the case of TRPV1, it is enriched in the filopodial tips^9, 22–24^. In case of TRPV4, rapid translocation of the GFP-tagged channels to the filopodial tips is observed after activation of TRPV4^22^. However, in the case of TRPV2, it is mainly located at the filopodial base and even after activation, it does not translocate to the filopodial tips in general, but mainly present in the plasma membrane. Interestingly, TRPV2 is known to translocate rapidly to other actin-based structures such as in the nascent phagosome (of macrophages) and podosome where it is involved in the podosome assembly^28, 63^. Such fine differences may indicate the subtle differences of TRP channels associations with different motor proteins present in the filopodia and lamellipodia^64^. Never-the-less, we demonstrate that TRPV2 physically interacts with submembranous cytoskeleton, mainly with actin cytoskeleton. This aspect matches well with the previous report that the C-terminus of TRPV4 interacts with soluble actin^22^. In this regard, it is important to note that the C-terminus of TRPV2 also interacts with tubulin^33, 63^. At present it is not known if and how TRPV2 regulates cytoskeleton and *vice versa*. Such interactions can be critical for membrane-stretch induced activation of TRPV2. Such interactions might also be important for the regulation of membrane ruffling, membrane spreading, filopodia formation and involvement of different kinase signaling events. However, further experiments are needed to dissect the signaling events in more details.

Study of vanilloid member such as TRPV2 and TRPV3 have been limited in the absence of specific pharmacological agonist and antagonist. But recent studies primarily have shown that some of these drugs such as Probeneid, Tranilast and 2APB are relatively specific on TRPV2. Our study is also limited in terms of the specific pharmacological activation and inhibition. To remove the doubt of non-specific activation of other protein we have used very low concentration of drugs which have been reported in previous studies. In conclusion, our work suggests that TRPV2 detects specific stimuli and influences the neurons to take decisions in the certain process of neuritogenesis such as in neurite initiation, neurite branching events which are other way considered as stochastic events. Such findings may help us to understand the cellular and molecular basis of different sensory functions and their implications in different pathophysiological conditions, such as in neurodegeneration.

## Acknowledgements

Funding from NISER, DST (Govt. India, grant number SR/SO/HS/0040/2012) and DBT (Govt. India, grant number BT-BRB-TF-2-2011 and BT/PR8004/MED/30/988/2013) are acknowledged. We thanks to Prof. E.D. Gundelfinger and Dr. KH Smalla (Leibniz Institute for Neurobiology, Magdeburg) for providing synaptosomal fraction. We acknowledge the support and critical reagents from Prof. Ferdinand Hucho (Freie Universität Berlin) and Dr. Ricarda Jahnel (Freie Universität Berlin). The funders had no role in study design, data collection and preparation of the manuscript. Imaging facility from NISER is appreciated.

## Author contributions and notes

CG conceived the idea. MY did all the experiments, prepared the figures and graphs. MY and CG analysed the data, and are involved in the MS preparation. The authors declare that the funding agencies have no role in the experimental design, data generation, data representation and decision to publish.

## References

1. Hutchins, B. I. Competitive outgrowth of neural processes arising from long-distance cAMP signaling. Sci Signal, 3, jc1 (2010)

2. Hutchins, B. I., & Kalil, K. Differential outgrowth of axons and their branches is regulated by localized calcium transients. J Neurosci, 28, 143–153 (2008)

3. Bedlack, R. S., Jr. Wei, M., & Loew, L. M. Localized membrane depolarizations and localized calcium influx during electric field-guided neurite growth. Neuron, 9, 393–403 (1992)

4. Polak, K. A., Edelman, A. M., Wasley, J. W., & Cohan, C. S. A novel calmodulin antagonist, CGS9343B, modulates calcium-dependent changes in neurite outgrowth and growth cone movements. J Neurosci, 11, 534–542 (1991)

5. Ramakers, G. J. et al The role of calcium signaling in early axonal and dendritic morphogenesis of rat cerebral cortex neurons under non-stimulated growth conditions. Brain Res Dev Brain Res, 126, 163–172 (2001)

6. Schindelholz, B., & Reber B. F. L-type Ca^2+^ channels and purinergic P2X2 cation channels participate in calcium-tyrosine kinase-mediated PC12 growth cone arrest. Eur J Neurosci, 12, 194–204 (2000)

7. Nishiyama, M. et al. Cyclic AMP/GMP-dependent modulation of Ca^2+^ channels sets the polarity of nerve growth-cone turning. Nature, 423, 990–995 (2003)

8. Goswami, C., Schmidt, H., & Hucho F. TRPV1 at nerve endings regulates growth cone morphology and movement through cytoskeleton reorganization. FEBS J, 274, 760–772 (2007)

9. Goswami, C., & Hucho, T. TRPV1 expression-dependent initiation and regulation of filopodia. J Neurochem, 103, 1319–1333 (2007)

10. Cohen M. R. et al. Nerve Growth Factor regulates Transient Receptor Potential Vanilloid 2 via extracellular Signal-Regulated Kinase signaling to enhance neurite outgrowth in developing neurons. Mol Cell Biol, 35, 4238–4252 (2015)

11. Sigerson, C. D., Dipollina, C. J., & Fornaro, M. Effects of chemical and physical cues in enhancing neuritogenesis and peripheral nerve regeneration. Neural Regen Res, 11, 220–221 (2016)

12. Gomez, T. M., & Spitzer, N. C. *In vivo* regulation of axon extension and path finding by growth-cone calcium transients. Nature 397, 350–355 (1999)

13. Oyama, K. et al. Triggering of high-speed neurite outgrowth using an optical microheater. Sci Rep, 5, 16611 (2015)

14. Pellegrino, M. & Pellegrini, M. Mechanosensitive channels in neurite outgrowth. Curr Top Membr. 59, 111–25 (2007)

15. Denis, V., & Cyert, M. S. Internal Ca(2+) release in yeast is triggered by hypertonic shock and mediated by a TRP channel homologue. J Cell Biol, 156, 29–34 (2002)

16. Zhou, X. L., Batiza, A. F., Loukin, S. H., Palmer, C. P., Kung, C., & Saimi, Y. The transient receptor potential channel on the yeast vacuole is mechanosensitive. Proc Natl Acad Sci U S A, 100, 7105–7110 (2003)

17. de Bono, M., Tobin, D. M., Davis, M. W., Avery, L., & Bargmann, C. I. Social feeding in *Caenorhabditis elegans* is induced by neurons that detect aversive stimuli. Nature, 419, 899–903 (2002)

18. Stowers, L., Holy, T. E., Meister, M., Dulac, C., & Koentges, G. Loss of sex discrimination and male-male aggression in mice deficient for TRP2. Science, 295, 1493–1500. (2002)

19. Zhang, Y. et al. Coding of sweet, bitter, and umami tastes: different receptor cells sharing similar signaling pathways. Cell, 112, 293–301. (2003)

20. Goswami, C., Dreger, M., Jahnel, R., Bogen, O., Gillen, C., & Hucho, F. Identification and characterization of a Ca^2+^-sensitive interaction of the vanilloid receptor TRPV1 with tubulin. J Neurochem. 91, 1092–103. (2004)

21. Goswami C. Structural and functional regulation of growth cone, filopodia and synaptic sites by TRPV1. Commun Integr Biol. 3, 614–8. (2010)

22. Goswami C., Kuhn J., Heppenstall P. A., & Hucho T. Importance of non-selective cation channel TRPV4 interaction with cytoskeleton and their reciprocal regulations in cultured cells. PLoS One, 5, e11654. (2010)

23. Goswami C., Rademacher N., Smalla K. H., Kalscheuer V., Ropers H. H., Gundelfinger E. D., & Hucho T. TRPV1 acts as a synaptic protein and regulates vesicle recycling. J Cell Sci, 123, 2045–2057. (2010)

24. Goswami, C. et al. Estrogen destabilizes microtubules through an ion-conductivity-independent TRPV1 pathway. J Neurochem. 117, 995–1008. (2011)

25. Greka, A, Navarro, B, Oancea, E, Duggan, A, & Clapham, D. E. TRPC5 is a regulator of hippocampal neurite length and growth conemorphology. Nat Neurosci. 6, 837–45. (2003)

26. Wang, G. X., Poo, M. M. Requirement of TRPC channels in netrin-1-induced chemotropic turning of nerve growth cones. Nature, 434, 898–904. (2005)

27. Sugio, S., Nagasawa, M., Kojima, I., Ishizaki, Y., Shibasaki, K. Transient receptor potential vanilloid 2 activation by focal mechanical stimulation requires interaction with the actin cytoskeleton and enhances growth cone motility. FASEB J. 31, 1368–1381. (2017)

28. Link, T. M., Park, U., Vonakis, B. M., Raben, D. M., Soloski, M. J. & Caterina, M. J. TRPV2 has a pivotal role in macrophage particle binding and phagocytosis. Nat Immunol, 11, 232–239. (2010)

29. Katanosaka, Y. et al. TRPV2 is critical for the maintenance of cardiac structure and function in mice. Nat Commun, 5, 3932. (2014)

30. Pavlov, V. A., & Tracey, K. J. Neural circuitry and immunity. Immunol Res, 63, 38–57. (2015)

31. Veiga-Fernandes, H., & Mucida, D. Neuro-Immune Interactions at Barrier Surfaces. Cell, 165(4), 801–811. (2016)

32. Chavan, S. S., Pavlov, V. A., & Tracey, K. J. Mechanisms and Therapeutic Relevance of Neuro-immune Communication. Immunity, 46, 927–942. (2017)

33. Jahnel, R. Investigations of molecular pain perception mechanisms, especially biochemical characterization of the thermosensitive vanilloid receptors TRPV1 and TRPV2. (http://www.diss.fu-berlin.de/diss/receive/FUDISS_thesis_000000001749) (2005)

34. Klarenbeek, J., Goedhart, J., van Batenburg, A., Groenewald, D. & Jalink, K. Fourth-generation epac-based FRET sensors for cAMP feature exceptional brightness, photostability and dynamic range: characterization of dedicated sensors for FLIM, for ratiometry and with high affinity. PLoS One 10, e0122513. (2015)

35. Montell, C. Exciting trips for TRPs. Nat Cell Biol, 6, 690–692. (2004)

36. Li, Y. et al. Essential role of TRPC channels in the guidance of nerve growth cones by brain-derived neurotrophic factor. Nature, 434, 894–898. (2005)

37. Shim, S. et al. XTRPC1-dependent chemotropic guidance of neuronal growth cones. Nat Neurosci, 8, 730–735. (2005)

38. Bender, F. L. et al. The temperature-sensitive ion channel TRPV2 is endogenously expressed and functional in the primary sensory cell line F-11. Cell Physiol Biochem, 15, 183–194. (2005)

39. Jahnel, R., Bender, O., Munter, L. M., Dreger, M., Gillen, C., & Hucho, F. Dual expression of mouse and rat VRL-1 in the dorsal root ganglion derived cell line F-11 and biochemical analysis of VRL-1 after heterologous expression. Eur J Biochem, 270, 4264–4271. (2003)

40. Mitchison, T. J., & Cramer, L. P. Actin-based cell motility and cell locomotion. Cell, 84, 371–379. (1996)

41. Small, J. V., & Resch, G. P. The comings and goings of actin: coupling protrusion and retraction in cell motility. Curr Opin Cell Biol, 17, 517–523. (2005)

42. Small, J. V., Stradal, T., Vignal, E., & Rottner, K. The lamellipodium: where motility begins. Trends Cell Biol, 12, 112–120. (2002)

43. Wood, W., & Martin, P. Structures in focus-filopodia. Int J Biochem Cell Biol, 34, 726–730. (2002)

44. da Silva, J. S., & Dotti, C. G. Breaking the neuronal sphere: regulation of the actin cytoskeleton in neuritogenesis. Nat Rev Neurosci, 3, 694–704. (2002)

45. Faix, J., & Rottner, K. The making of filopodia. Curr Opin Cell Biol, 18, 18–25. (2006)

46. Dehmelt, L., & Halpain, S. Actin and microtubules in neurite initiation: are MAPs the missing link? J Neurobiol, 58, 18–33. (2004)

47. Sainath, R., & Gallo, G. Cytoskeletal and signaling mechanisms of neurite formation. Cell Tissue Res, 359, 267–278. (2015)

48. Kuner, R. (2010) Central mechanisms of pathological pain. Nat Med, 16, 1258–1266.

49. Neumann, S., Doubell, T. P., Leslie, T., & Woolf, C. J. Inflammatory pain hypersensitivity mediated by phenotypic switch in myelinated primary sensory neurons. Nature, 384, 360–364. (1996)

50. Zhou, N et al. Suppression of KV7/KCNQ potassium channel enhances neuronal differentiation of PC12 cells. Neuroscience. 333, 356–67. (2016)

51. Yamashita, N. et al. Voltage-gated calcium and sodium channels mediate Sema3A retrograde signaling that regulates dendritic development. Brain Res. 1631, 127–36. (2016)

52. Shibasaki, K., Murayama, N., Ono, K., Ishizaki, Y., Tominaga, M. TRPV2 enhances axon outgrowth through its activation by membrane stretch in developing sensory and motor neurons. J Neurosci, 30, 4601–4612. (2010)

53. Nedungadi, T. P., Dutta, M., Bathina, C. S., Caterina, M. J., & Cunningham, J. T. Expression and distribution of TRPV2 in rat brain. Exp Neurol, 237, 223–237. (2012)

54. Bang, S., Kim, K. Y., Yoo, S., Lee, S. H., & Hwang, S. W. Transient receptor potential V2 expressed in sensory neurons is activated by Probenecid. Neurosci Lett, 425, 120–125. (2007)

55. Yamamoto, Y., Sato Y., & Taniguchi, K. Distribution of TRPV1- and TRPV2-immunoreactive afferent nerve endings in rat trachea. J Anat, 211, 775–783. (2007)

56. Zhang, H., Xiao, J., Hu, Z., Xie, M., Wang, W., & He, D. Blocking transient receptor potential vanilloid 2 channel in astrocytes enhances astrocyte-mediated neuroprotection after oxygen-glucose deprivation and reoxygenation. Eur J Neurosci, 44, 2493–2503. (2016)

57. Jang, Y. et al. Axonal neuropathy-associated TRPV4 regulates neurotrophic factor-derived axonal growth. J Biol Chem, 287, 6014–6024. (2012)

58. Park, U., Vastani, N., Guan, Y., Raja, S. N., Koltzenburg, M., & Caterina, M. J. TRP vanilloid 2 knock-out mice are susceptible to perinatal lethality but display normal thermal and mechanical nociception. J Neurosci, 31, 11425–11436. (2011)

59. Anava, S., Greenbaum, A., Ben Jacob, E., Hanein, Y., & Ayali, A. The regulative role of neurite mechanical tension in network development. Biophys J, 96, 1661–1670. (2009)

60. Bornschlögl, T. & Bassereau, P. The sense is in the fingertips: The distal end controls filopodial mechanics and dynamics in response to external stimuli. Commun Integr Biol. 6, e27341. (2013)

61. Xiong, Y., Lee, A. C., Suter, D. M., & Lee, G. U. Topography and nanomechanics of live neuronal growth cones analyzed by atomic force microscopy. Biophys J, 96, 5060–5072. (2009)

62. Heidemann, S. R., & Buxbaum, R. E. Mechanical tension as a regulator of axonal development. Neurotoxicology 15, 95–107. (1994)

63. Nagasawa, M., & Kojima, I. Translocation of calcium-permeable TRPV2 channel to the podosome: Its role in the regulation of podosome assembly. Cell Calcium, 51, 186–193. (2012)

64. Majhi, R., Sardar, P., Goswami, L., & Goswami, C. Right time - right location - right move: TRPs find motors for common functions. Channels (Austin). 5, 375–81. (2011)

65. Goswami, C. Identification of tubulin as a TRPV1-interacting protein and functional characterization of TRPV1-cytoskeleton regulation. http://www.diss.fu-berlin.de/diss/receive/FUDISS_thesis_000000002142 (2006)

